# Extracting thermodynamic properties from van ’t Hoff plots with emphasis on temperature-sensing ion channels

**DOI:** 10.1101/2023.06.02.543442

**Authors:** Jakob Tómas Bullerjahn, Sonya M. Hanson

## Abstract

Transient receptor potential (TRP) ion channels are among the most well-studied classes of temperature-sensing molecules. Yet, the molecular mechanism and thermodynamic basis for the temperature sensitivity of TRP channels remains to this day poorly understood. One hypothesis is that the temperature-sensing mechanism can simply be described by a difference in heat capacity between the closed and open channel states. While such a two-state model may be simplistic it nonetheless has descriptive value, in the sense that it can be used to to compare overall temperature sensitivity between different channels and mutants. Here, we introduce a mathematical framework based on the two-state model to reliably extract temperature-dependent thermodynamic potentials and heat capacities from measurements of equilibrium constants at different temperatures. Our framework is implemented in an open-source data analysis package that provides a straightforward way to fit both linear and nonlinear van ‘t Hoff plots, thus avoiding some of the previous, potentially erroneous, assumptions when extracting thermodynamic variables from TRP channel electrophysiology data.

## I. INTRODUCTION

An organism’s ability to sense its environment is crucial to its survival. One of the most well-studied families of biological temperature sensors in humans and other eukaryotes is the transient receptor potential (TRP) family of ion channels [1]. Members of this family have temperature sensitivity across the biologically relevant range of temperatures, but the most well-known are the heat and capsaicin-sensitive TRPV1 [2] in the TRPV subfamily, and the cold and menthol sensitive TRPM8 [3, 4] in the TRPM subfamily. Hypotheses about the principles guiding the temperature-sensitivity of TRP channels were already being postulated within a few years of their discovery, with proposed mechanisms relating to phenomena from voltage-sensing to elongations of open channel burst times [5, 6]. However, while these molecules have been identified as intrinsically sensitive to temperature [7] and playing a critical role as temperature sensors in our nervous system [8–10], we still do not understand the molecular and thermodynamic mechanism(s) that dictates their temperature-dependent activation.

One characteristic of TRP ion channels that seems clear are the large positive enthalpy differences between states for heat-sensitive TRPs like TRPV1 [5, 11] and large negative enthalpy differences for cold-sensitive TRPs like TRPM8 [11, 12]. Entropy and enthalpy differences between the open and closed states of a channel can be extracted from linear fits to the logarithm of the equilibrium constant *K*_eq_ as a function of the reciprocal temperature 1*/T* if said thermodynamic potentials are independent of temperature. However, as is well known in the literature of physical biochemistry, large conformational changes in proteins are usually accompanied by changes in their heat capacities, which leads to temperature-dependent enthalpies and entropies [13]. This is the premise of a modelfree framework proposed by Clapham and Miller [11], which can explain both cold and heat-sensitive changes in the equilibrium constant. While it remains debated whether temperature-dependent gating in channels is also accompanied by observable changes in heat capacity [14], this is the main mechanism to induce temperature dependence in the relevant thermodynamic potentials.

Here, we embrace the approach of Clapham and Miller [11], and introduce a procedure to reliably extract temperature-dependent thermodynamic potentials and heat capacities from equilibrium constant measurements performed at different temperatures. We thereby assume that a TRP channel can, to a first approximation, be described as a two-state system, which may not provide the same mechanistic insight as more involved models [15], but has the benefit of being universally applicable and allows for a direct comparison of thermodynamic variables obtained for different ion channels or the same channel at differing experimental conditions. Our theory is implemented in an open-source data analysis package [16] written in Julia [17], and should provide practitioners a straightforward way to fit linear and nonlinear van ‘t Hoff plots, thus avoiding previous potentially false assumptions about the nature of temperature sensors.

The paper is structured as follows. In Sec. II A, we list the thermodynamic relations relevant to our discussion, and briefly review their common use in the literature of temperature sensors. Section II B introduces cubic splines as continuously differentiable functions used to fit discrete measurements of ln(*K*_eq_). Under the assumption of a two-state model, the differentiability of splines allows us to calculate robust estimates for the thermodynamic potentials Δ*H*(*T*) and Δ*S*(*T*), and the associated heat capacity difference Δ*C*_*p*_(*T*). To avoid overfitting, we rely on a Bayesian information criterion (BIC) [18] to penalize splines with many degrees of freedom, as described in Sec. II C. For illustrative purposes, we apply our data analysis package to two distinct data sets in Sec. III, and conclude in Sec. IV with a summary of our results.

## II. THEORY

One of the simplest ways to model a TRP channel is to treat it as a two-state system. Every channel in an ensemble of channels can then either be in the open or closed state, such that the composition of the ensemble is encoded in the equilibrium constant *K*_eq_:

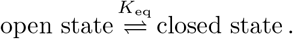

In electrophysiological experiments the charge current through a single channel or a collection of channels is measured at different temperatures, which can be used to calculate the so-called “open probability” *P*, i.e., the probability of finding a channel in the open state. The equilibrium constant *K*_eq_ and the open probability *P* are related via

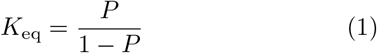

for a two-state system. Note that macroscopic ionic currents are subject to a multitude of additional sources of variability that can predominate at the temperature extremes, where channel activity is either very low or near-maximal. Because the quality of the associated *P* and *K*_eq_ estimates is directly affected, we recommend users to carefully select the temperature range of the data to be fitted to avoid contributions from sources of signal variability unrelated to channel gating.

### A. Thermodynamic description

Thermodynamics tells us, on the one hand, that the differentials of enthalpy *H* and entropy *S* are related via

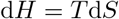

for systems at constant pressure, where *T* denotes the absolute temperature. On the other hand, it can be shown that the heat capacity *C*_*p*_ at constant pressure satisfies

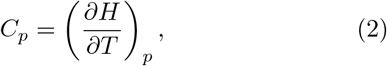

where *∂/∂T* denotes a partial derivative with respect to *T* and the index of the bracket reveals which quantity is being held constant (in this case it is the pressure *p*). We therefore conclude that the enthalpy and entropy differences between two metastable states must be integral functions of the heat capacity difference 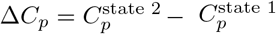 between the states, i.e.,

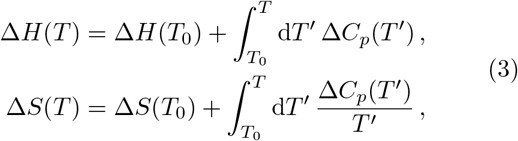

for some arbitrary reference temperature *T*_0_. Note that Δ*H*(*T*) and Δ*S*(*T*) are independent of *T* whenever Δ*C*_*p*_(*T*) = 0, e.g., for bistable systems whose states have the same heat capacity. However, in the case of protein folding, we know that large heat capacity differences exist between their folded and unfolded state [13].

Another thermodynamic potential of interest is the Gibbs free energy, which is given by

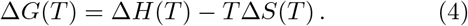

It can be related to the equilibrium constant *K*_eq_ of the two-state system via the fundamental relation of chemical thermodynamics, namely

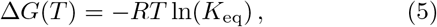

where *R* = 8.314 462 618 153 24 J mol^−1^ K^−1^ denotes the molar gas constant. The logarithm of the equilibrium constant and its derivative with respect to *T* therefore have the form

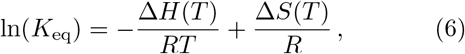

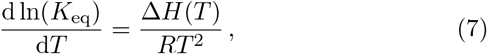

where the latter is the well-known van ‘t Hoff equation, which is sometimes also written as follows:

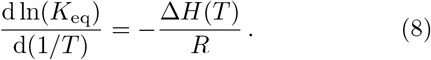

Equations (6) and (8) reveal that a so-called van ‘t Hoff plot, where ln(*K*_eq_) is plotted against the reciprocal of the absolute temperature *T*, will be linear whenever Δ*H* and Δ*S* are constant with respect to temperature. The thermodynamic potentials can then be read off the slope and intercept of ln(*K*_eq_), respectively. This convenient fact often seems to guide the decision of practitioners to fit their data to straight lines, even when the van ‘t Hoff plot is highly nonlinear (see, e.g., Refs. 12 and 19), which can be an indication for temperature-dependent behavior. Overall, it is important to note that Eqs. (7) and (8) are valid for all functions Δ*H*(*T*) and Δ*S*(*T*) of the form given in Eq. (3), and not just constant thermodynamic potentials.

A popular empirical measure of temperature sensitivity is *Q*_10_, which is used to characterize temperature sensitivity in electrophysiological experiments on TRP channels [20, 21]. It is defined as the ratio of *K*_eq_ measured at two temperatures that are 10 K apart, i.e.,

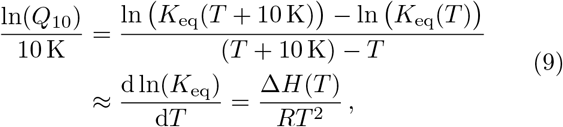

The reason for its wide-spread use is the fact that on a logarithmic scale it approximately reproduces the van ‘t Hoff equation, i.e.,

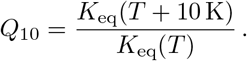

and can therefore be used to estimate Δ*H*(*T*). Again, it is common practice to assume that Δ*H* is temperature-independent and ln(*Q*_10_) is therefore only evaluated at a single temperature *T*, which can lead to arbitrary and skewed results whenever Δ*H*(*T*) = const. is not satisfied.

Here, we call for a different approach to extract thermodynamic information from measured *K*_eq_-values, without any ad-hoc assumptions. Instead of performing a linear fit of ln(*K*_eq_) plotted against *T* ^−1^, we instead propose to extract Δ*H*(*T*), Δ*S*(*T*) and Δ*G*(*T*) from Eqs. (4), (5) and (7) [or, equivalently, Eq. (8)], as discussed in Sec. II B. In cases, where Δ*H*(*T*) and Δ*S*(*T*) vary strongly with the temperature, this novel approach allows also us to estimate the change in heat capacity Δ*C*_*p*_.

### B. Spline fitting of discrete data points

Extracting derivatives from discrete points, e.g., via finite differences, can be somewhat tricky, because direct numerical differentiation amplifies the noise in the data. Therefore, advanced methods like spline-fitting [22] should be considered to construct continuous curves for ln(*K*_eq_) and its derivatives.

Consider a piecewise continuous function

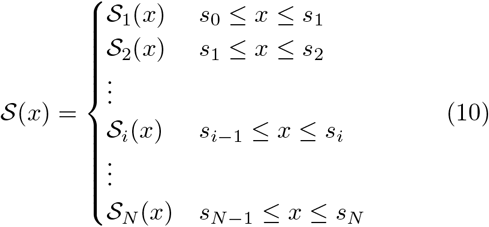

made up of third-order polynomials of the form

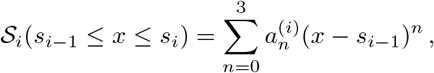

which are joined together in the spline knots *s*_*i*_. The function 𝒮 (*x*) is known as a cubic spline and satisfies the continuity conditions

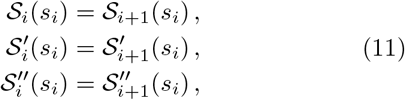

Where the notation *S*^*l*^(*x*) = d*S*(*x*)*/*d*x* and *S*^*ll*^(*x*) = d^2^*S*(*x*)*/*d*x*^2^ was introduced to abbreviate the expressions. We also require some appropriate boundary conditions, e.g., so-called natural boundary conditions given by

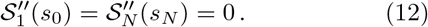

Equations (11) and (12) constrain the values of the spline coefficients 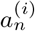, such that only *N* + 1 of them can be varied independently.

In our fitting procedure, the edge knots *s*_0_ and *s*_*N*_ are held fixed, while the “inner” knots *{s*_1_, …, *s*_*N*−1_*}* are allowed to vary within the interval [*s*_0_, *s*_*N*_]. We also vary the values 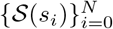 of the splines at the knots (see Fig. 1). The best fit of 𝒮 (*x*) to the data 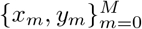 minimizes the sum of squared residuals between the data points and the spline, i.e.,

**Figure 1.**
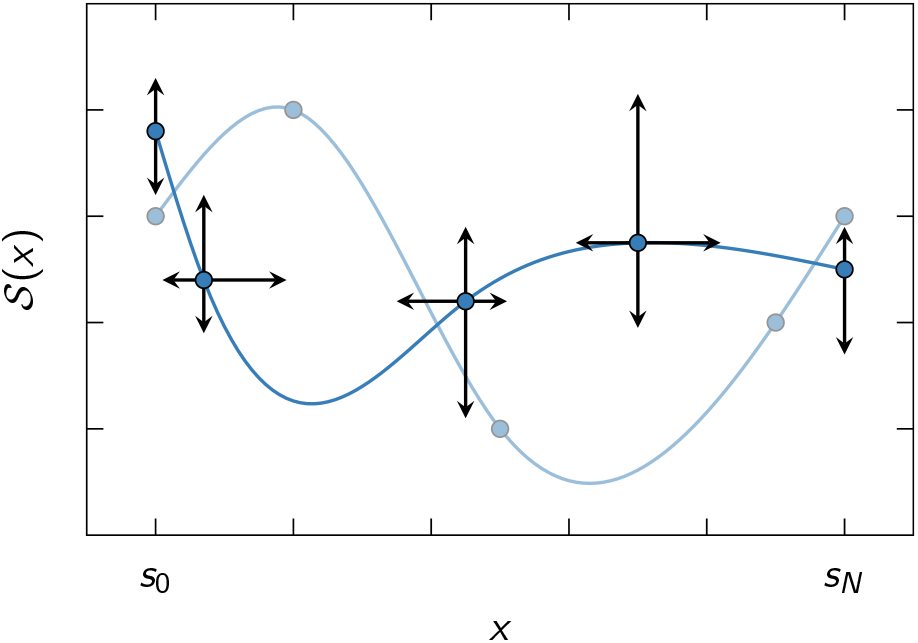
Visualizing the principle of spline fitting. When fitting a cubic spline 𝒮(*x*) to some data points 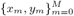, we exploit the fact that for every set of knot coordinates (*s*_0_, 𝒮 (*s*_0_)), (*s*_1_, 𝒮 (*s*_1_)), …, (*s*_*N*_, 𝒮 (*s*_*N*_)) (blue circles) there exists a unique cubic spline (blue solid lines) satisfying the boundary conditions in Eq. (12). Thus, by varying the knot coordinates (black arrows), we can change the shape of the spline to minimize *χ*^2^ in Eq. (13). Note that the edge knots at *x* = *s*_0_ and *x* = *s*_*N*_ can only be varied in *y*-direction, whereas the “inner” knots are allowed to take arbitrary *x*-values within the interval [*s*_0_, *s*_*N*_].

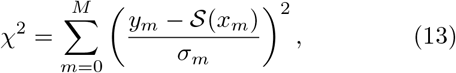

where *σ*_*m*_ denotes the standard error of *y*_*m*_. Here, we consider *y*_*m*_ = ln *K*_eq_(*x*_*m*_) for either a linear (*x*_*m*_ = *T*_*m*_) or reciprocal temperature scale (*x*_*m*_ = 1*/T*_*m*_), and set the values of *s*_0_ and *s*_*N*_ equal to the lowest and highest values of *x*_*m*_ found in the data set, respectively. The reason why we consider both scales is because one cannot distinguish between a linear and reciprocal temperature dependence for the temperature ranges realized in electrophysiological experiments (see also Fig. 2).

Evaluating 𝒮*x*(*T*) for the parameters that minimize Eq. (13) therefore gives the best estimate of ln (*K*_eq_(*T*)), which can be used to extract the heat capacity difference Δ*C*_*p*_(*T*) and the thermodynamic potentials Δ*G*(*T*),

Δ*H*(*T*), and Δ*S*(*T*) as follows:

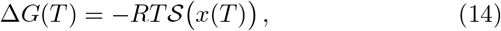

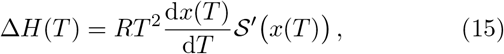

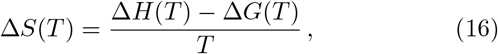

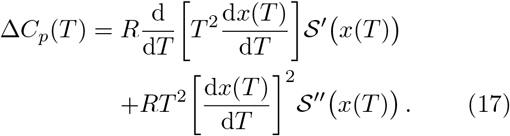

Note that for *x* = *T* we have d*x/*d*T* = 1, whereas the reciprocal relation *x* = 1*/T* gives d*x/*d*T* = −*T* ^−2^ for which the first term of Eq. (17) vanishes.

### C. Model selection

The choice between a linear and reciprocal fit, as well as the number of spline knots *N* + 1, gives rise to a multitude of models that fit the data set to varying degree. For a fixed *N*, one can distinguish between the qualities of a linear and a reciprocal fit by comparing their corresponding *χ*^2^ values, but if *N* is allowed to vary then models with *N ≫* 1 will always be preferred. We therefore propose the use of an information criterion [18] to penalize models with too many fit parameters. By interpreting Eq. (13) as a negative log-likelihood for Gaussian distributed residuals, we obtain the following BIC:

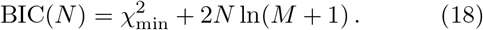

Equation (18) is evaluated using the optimal values for the 2*N* spline parameters 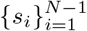 and 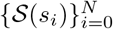 that minimize *χ*^2^ [Eq. (13)], resulting in the minimum value 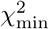. The model that best fits the data, while avoiding overfitting, minimizes BIC(*N*) with respect to *N*.

Our data analysis package automatically varies *N*, finds the associated optimal parameter values that minimize *χ*^2^, and subsequently calculates the corresponding BIC-value. It finally returns the model and associated parameter values that best fit the data at hand.

### D. Extension to multi-state models

In principle, our data-fitting approach can be extended to (and used to generalize) models with multiple states, such as the ones presented in Ref. 15, by replacing Eq. (1) with an expression *P* (*T*) for the open probability involving multiple spline functions 𝒮 ^(*k*)^ *x*((*T*)) with *k* = 1, 2, …. The fitting must then be performed on the level of *P*, instead of ln(*K*_eq_), which implies that Eq. (13) gets replaced with

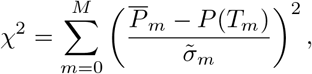

where 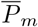 and 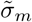 are the mean and associated standard error of the measured open probability at temperature *T*_*m*_. The corresponding BIC takes the form

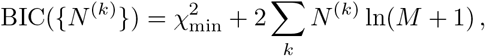

where *N* ^(*k*)^ + 1 is the number of spline knots in 𝒮 ^(*k*)^(*x*).

For a concrete example, consider the four-state model in Ref. 15, where the open probability is given by

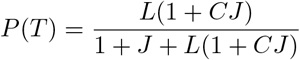

and the coefficients *L, C*, and *J* are related to the equilibrium constants between the two open (“O”) and two closed (“C”) states as follows:

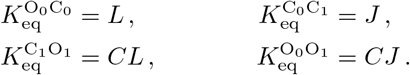

This model can be generalized by replacing *L, C*, and *J* with exp 𝒮 ^(*k*)^(*x*) |_*k*=1,2,3_, respectively, if all equilibrium constants are assumed to be temperature dependent. After fitting the data in analogy to the two-state case, Eqs. (14) to (17) can then be evaluated by replacing 𝒮 (*x*) with 𝒮 ^(1)^(*x*), 𝒮 ^(3)^(*x*), 𝒮 ^(1)^(*x*) + 𝒮 ^(2)^(*x*), and 𝒮 ^(2)^(*x*)+ 𝒮 ^(3)^(*x*) to extract the thermodynamic potentials and heat capacity differences related to all the different equilibrium constants.

Even though the generalization to multiple states is fairly straight-forward, our data analysis package currently only supports a two-state description.

## III. RESULTS AND DISCUSSION

For illustrative purposes, we applied the data analysis package to two previously published data sets, one for the warm-sensitive TRPV3 channel [19], and another for the heat and capsaicin-sensitive TRPV1 channel [5]. Each data set was analyzed by performing a van ‘t Hoff fit of measured values of ln(*K*_eq_) for different temperatures *T* to extract heat capacity differences and thermodynamic potentials, as described in Secs. II B and II C. Here, we deliberately avoid a direct comparison with the results of the associated publications, as it is not our intention to question their conclusions, but to demonstrate how our data analysis package works in practice.

In the case of the TRPV3 channel, we considered the measured open probabilities *p* that are tabulated in the source data associated with the extended data Fig. 1 in Ref. 19. For each temperature *T*_*m*_, we calculated the sample mean and variance of *P*_*m*_, i.e.,

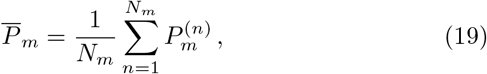

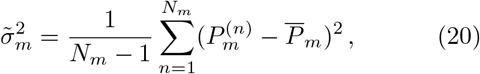

where 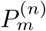 denotes the *n*th measurement (of *N*_*m*_ in total) of the open probability *P*_*m*_ at temperature *T*_*m*_. The equilibrium constant *K*_eq_ can be calculated via Eq. (1) and according to the variance formula of error propagation one has

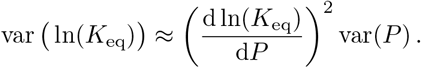

The data points and standard errors entering Eq. (13) are therefore given by

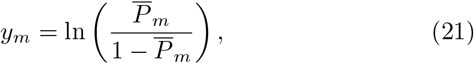

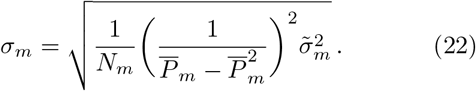

Our results for the TRPV3 data are shown in Fig. 3. The model that best fits the data is reciprocal in the temperature (*x* = 1*/T*) and contains *N* + 1 = 3 spline knots. The model predicts a temperature-dependent heat capacity difference Δ*C*_*p*_(*T*) that decreases monotonically beyond *T* ≈ 300 K [Fig. 3(c)]. The resulting enthalpy and entropy-temperature product differences, Δ*H*(*T*) and *T* Δ*S*(*T*), are therefore nonconstant and vary between −100 and +300 kJ mol^−1^ [Fig. 3(b)]. However, they mostly cancel each other out and give rise to a moderate free-energy difference, as seen in Fig. 3(a).

Figure 4 displays our results for the TRPV1 channel, where the data points *{T*_*m*_, *y*_*m*_*}* and standard errors *σ*_*m*_ were read off Fig. 2c of the original publication. The model that best fits the data is reciprocal in the temperature (*x* = 1*/T*) and has no inner knots, i.e. *N* + 1 = 2. At first this may seem somewhat surprising, considering the fact that the data are not perfectly linear in *T* ^−1^, but is essentially a good example of how our data analysis package avoids overfitting. Apparently, one does not gain sufficiently large improvements in the 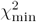 term of Eq. (18) to warrant a more complex model than one with Δ*C*_*p*_ = 0 and therefore constant thermodynamic potentials Δ*H* and Δ*S*. While we are here only illustrating the use of our data analysis package and want to refrain from making scientific assessment of the results at this time, we would like to make note of the narrow temperature range in this particular data set, so as the readers do not conclude that we are definitively claiming that TRPV1 has a vanishing Δ*C*_*p*_ across the physiological temperature range.

**Figure 2.**
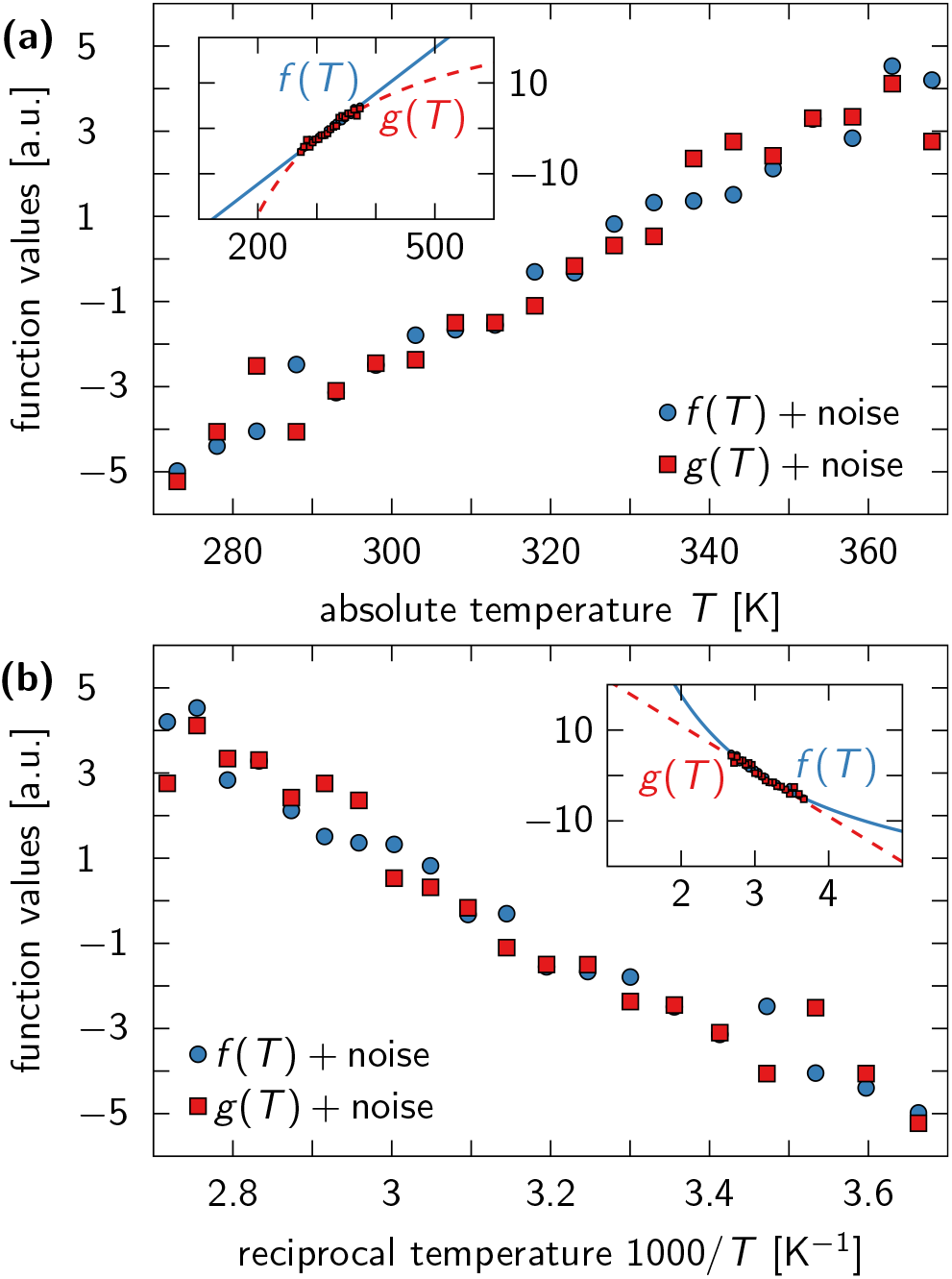
Distinguishing between linear and reciprocal functions on physiologically relevant temperature scales is impossible. (a) A linear function *f* (*T*) = *a* + *bT* (blue circles), perturbed by small noise, is plotted next to a reciprocal noisy function *g*(*T*) = *c* − *d/T* (red squares) on a temperature scale ranging from 0 °C to 100 °C. Both functions appear linear, because the absolute temperature is not varied by orders of magnitude to reveal the nonlinearity of *g*(*T*). (b) Same data as in (a) plotted on a reciprocal temperature scale. Again, both functions seem linear in 1*/T*, although only *g*(*T*) truly is. *Insets:* Same data as in (a) and (b) plotted on a wider temperature scale to visualize the linear and reciprocal trends of *f* (*T*) and *g*(*T*), respectively.

We can now compare our results to the output of alternative data analysis methods, such as the thermal coefficient *Q*_10_. In Fig. 5 we plot the enthalpies of Figs. 3(b) and 4(b) next to predictions that arise when Eq. (9) is solved for Δ*H*(*T*). The latter was evaluated using ln(*Q*_10_) = 𝒮 (*T* + 10 K) −𝒮 (*T*), where 𝒮 (*T*) ≡ 𝒮 (*x* = 1*/T*) because both data sets were fitted via reciprocal models. Figure 5(b) demonstrates that *Q*_10_ gives a decent estimate for the enthalpy whenever Δ*H* is independent of temperature. If this is not the case [Fig.5(a)], then the differences can become arbitrarily large, as can be seen in Fig. 5(a). Note that the discrepancy between the spline-fitting estimate and the *Q*_10_-estimate for Δ*H*(*T*) vanishes when the temperature difference entering the definition of *Q*_10_ goes to zero, i.e., when the finite-difference approximation in Eq. (9) becomes exact.

**Figure 3.**
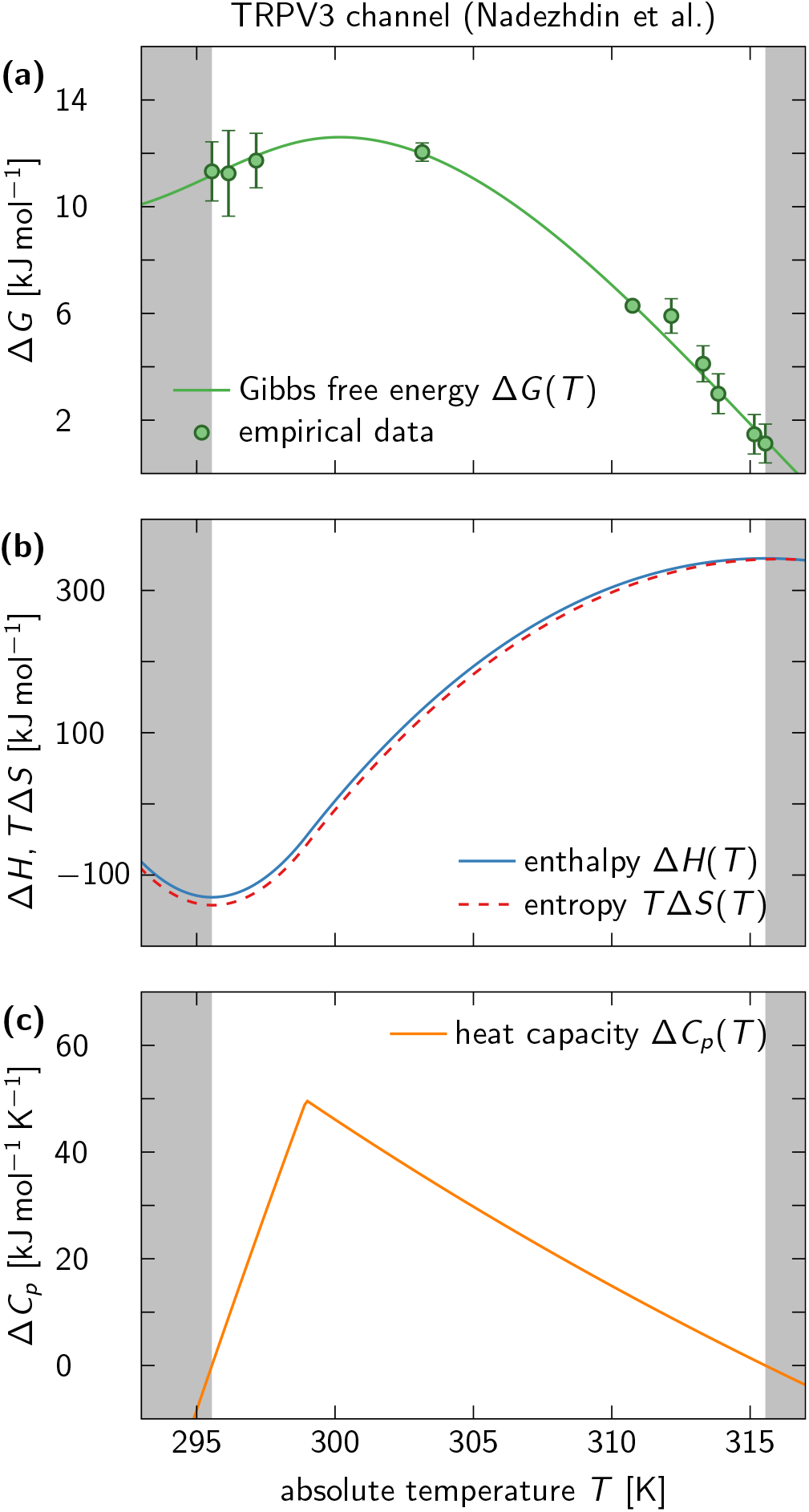
Thermodynamic potentials predicted from a van ‘t Hoff fit of TRPV3 channel data. The data are best fitted by a model that is reciprocal in the temperature with 3 spline knots. (a) Gibbs free energy Δ*G* as a function of *T*, calculated from data (points) and compared to model prediction (solid line). Shaded areas (gray) mark temperature intervals, where the trend of the cubic spline is no longer constrained by data points and can therefore not be trusted. (b) Enthalpy difference Δ*H* (blue solid line) and entropy-temperature product *T* Δ*S* (red dashed line) as functions of *T*. (c) Heat capacity difference Δ*C*_*p*_ as function of *T*. The temperature-dependence of Δ*H* and Δ*S* emerges from a nonzero Δ*C*_*p*_ predicted by the bestfitting model.

**Figure 4.**
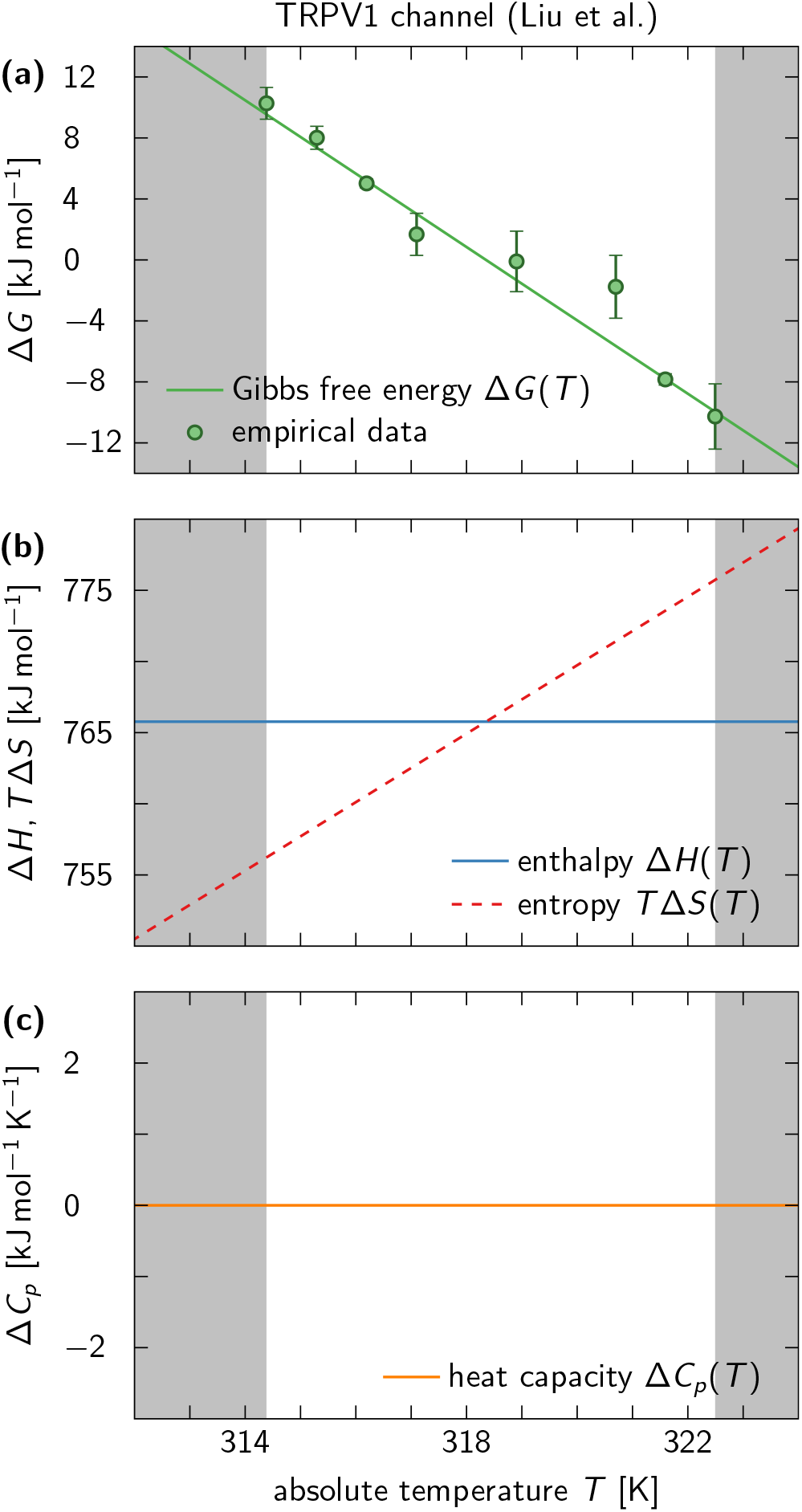
Thermodynamic potentials predicted from a van ‘t Hoff fit of TRPV1 channel data. The data are best fitted by a reciprocal model with 2 spline knots. (a-c) Same as in Fig. 3, except that here a model is preferred with vanishing heat capacity difference Δ*C*_*p*_, which results in Δ*H*, Δ*S* = const.

## IV. CONCLUSIONS

We have developed an open-access data analysis package [16] to reliably extract thermodynamic potentials and heat capacities from empirical measurements of equilibrium constants at different temperatures. Our package accounts for the fact that on physiologically relevant temperature scales one cannot distinguish between a linear and reciprocal temperature dependence (see Fig. 2), and therefore fits multiple models to the data, which vary in complexity (characterized by the number of parameters) and in the way they scale with temperature. A Bayesian information criterion [Eq. (18)] is used to select the model that best fits the data, while minimizing the number of model parameters to avoid overfitting. Our software can therefore be used to fit nonlinear van ‘t Hoff plots without any ad hoc assumptions and outperforms conventional methods, such as the thermal coefficient *Q*_10_ (see Fig. 5). Yet, we urge users to practice caution and not use our package to analyze data containing artifacts or unreasonably small error bars, because these can affect the resulting model selection and lead to faulty conclusions.

**Figure 5.**
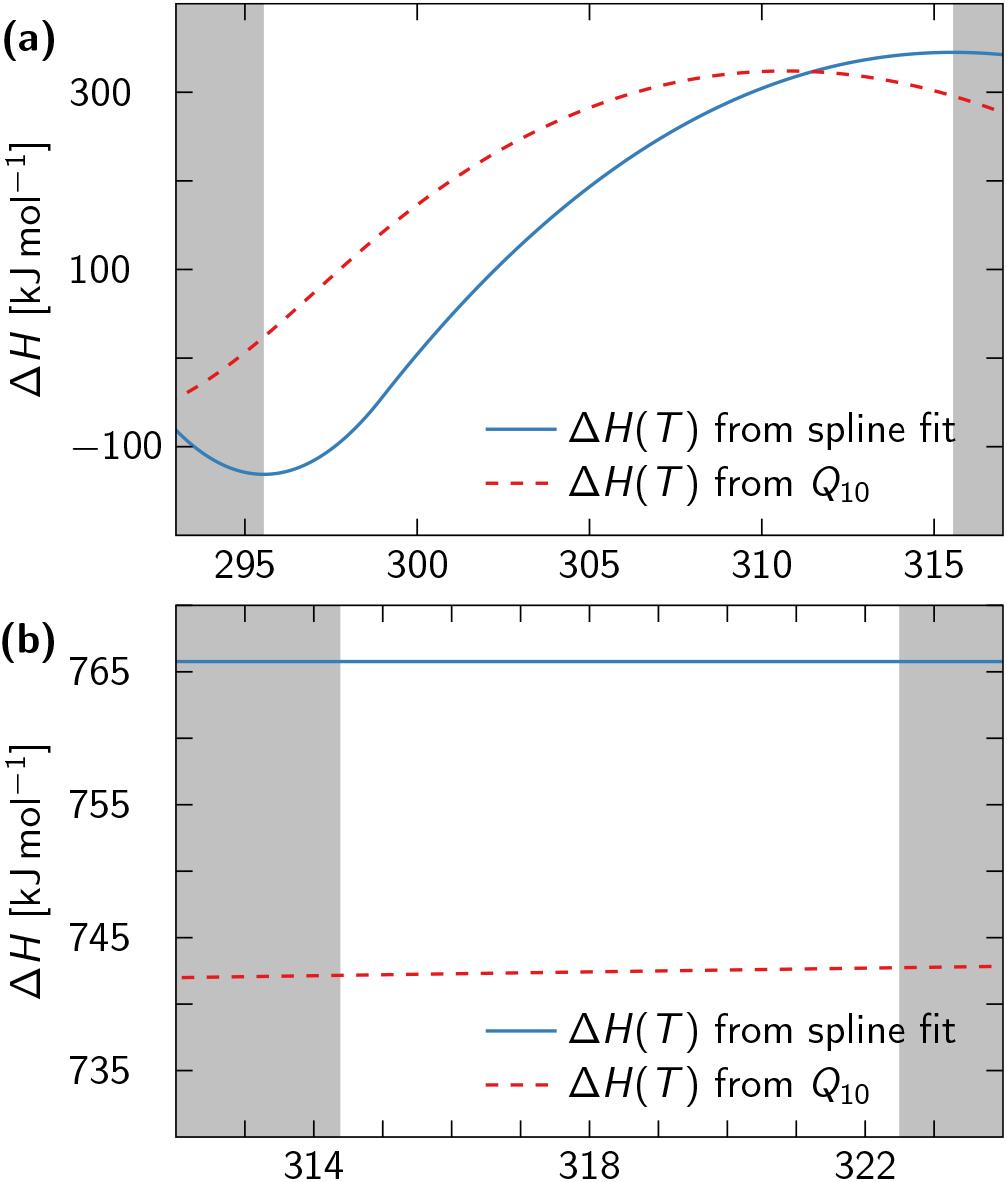
Comparison of enthalpy estimates obtained from van ‘t Hoff fits (blue solid lines) and *Q*_10_-based analysis (red dashed lines). (a) For the TRPV3 channel data analyzed in Fig. 3, the finite-difference approximation of the derivative with respect to *T* in Eq. (9) results in a vastly different Δ*H*(*T*) estimate than obtained from our spline-fitting procedure. (b) The van ‘t Hoff fit of the TRPV1 channel data predicted a temperature-independent enthalpy, for which *Q*_10_ provides a decent estimate of Δ*H* (in this case only 3% off).

To demonstrate the use of the data analysis package, we applied it to measurements of equilibrium constants for the temperature-sensitive TRPV1 and TRPV3 channels, respectively. For both data sets, we found that models with the functional form ln(*K*_eq_) = *f* (1*/T*), i.e., reciprocal in the temperature, were best suited to fit the data at hand, albeit with differing complexity. While the fit to the TRPV1 data predicted Δ*C*_*p*_ = 0, and therefore a constant enthalpy and entropy (see Fig. 4), an initially increasing and then monotonically decreasing Δ*C*_*p*_(*T*) was required to capture the extremely nonlinear trends seen in the TRPV3 data (Fig. 3). Note that our analysis relies on the common assumption that the TRPV1 and TRPV3 channels can, to a first approximation, be described as two-state systems. Even if this assumption is unlikely to hold for most (if not all) TRP channels, the thermodynamic variables that can be extracted from our approach provide a more reliable and accurate description of the temperature sensitivity of TRP channels than the results of a *Q*_10_-analysis or linear fits to a van ‘t Hoff plot. In principle, a more intricate analysis involving multiple states can also be conducted (see Sec. II D), but has not been implemented in our data analysis package for the simple reason that every mechanistic model would have to be implemented separately.

It is our belief that our data analysis tool will not only benefit the community of electrophysiologists studying temperature-sensitive channels, but also help researchers in chemistry and biochemistry to rigorously analyze their van ‘t Hoff plots. Temperature-dependent enthalpies and entropies open up exciting new possibilities in the theoretical modeling of the kinetics and dynamics of thermoresponsive systems, as the associated transition rates between the open and closed state intuitively must exhibit non-Arrhenius behavior. Whether such generalized models are applicable to TRP channels should be addressed in future research.

## ACKNOWLEDGMENTS

We thank Dr. Andrés Jara-Oseguera for fruitful discussions and critical comments on the manuscript. JTB acknowledges funding from the Max Planck society. The Flatiron Institute is a division of the Simons Foundation.

